## Supplemental Figures for "Fungal melanin suppresses airway epithelial chemokine secretion through blockade of calcium fluxing"

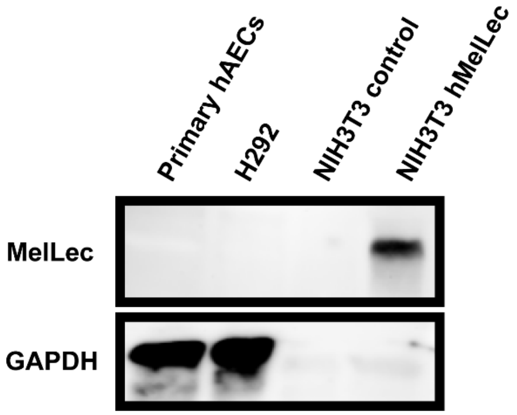


**Supplemental Figure 1. Primary hAECs and H292 cells lack MelLec receptor.** Human MelLec (hMelLec) protein expression in primary hAECs, H292 cells, or NIH3T3 fibroblast cells with or without hMelLec transfected into the cells. To ensure that there was not a low level of MelLec in epithelial cells, we added more protein lysate to primary hAECs and H292 when compared to NIH3T3 and NIH3T3 hMelLec cells. GAPDH was used to demonstrate protein loading.

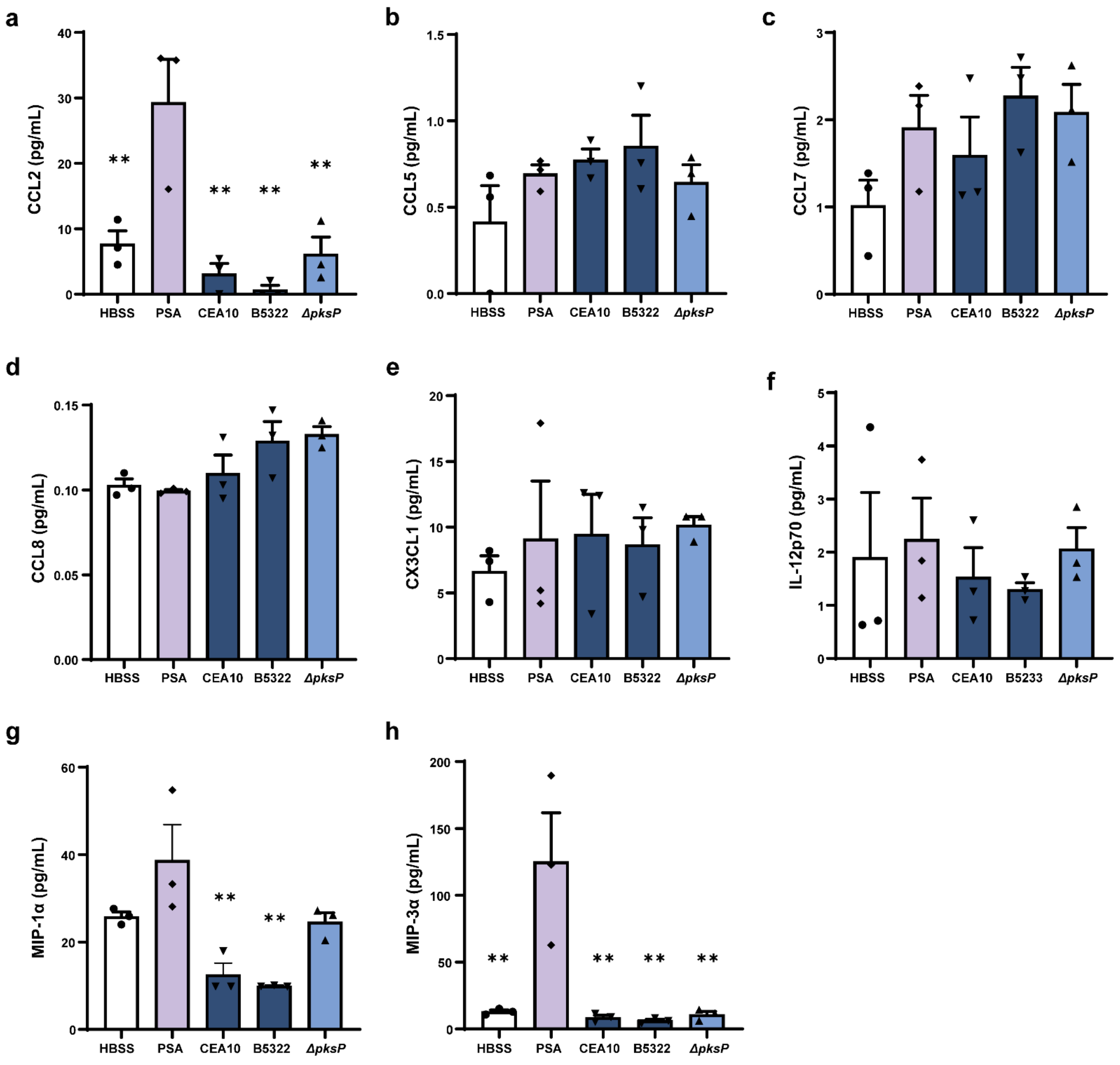


**Supplemental Figure 2. Multiplex assay revealed no difference in the secretion of other cytokine and chemokine in the presence of melanized *A. fumigatus*. (a)** CCL2, **(b)** CCL5, **(c)** CCL7, **(d)** CCL8, **(e)** CX3CL1, **(f)** IL12p70, **(g)** MIP-1α, and **(h)** MIP-3α in apical cell supernatants as measured by Luminex assay. Primary hAECs following 4h stimulation by media alone (HBSS), *P. aeruginosa* PAO1 strain (PSA), two different wildtype (WT) *A. fumigatus* strains (1 x $10^7/cm^2$), or *ΔpksP A. fumigatus* conidia (1 x $10^7/cm^2$). n=3. Data are represented as mean ± SEM. One-way ANOVA with Tukey's multiple comparisons test; **p<0.01 compared to PSA alone. The following cytokines/chemokines were tested but below the detection limit: CCL11, CCL13, CCL17, CCL19, CCL22, CCL24, CXCL1, CXCL5, CXCL9, CXCL10, CXCL11, CXCL13, GM-CSF, IFN-α, IL-1α, IL-1β, IL-11, IL-12p40, IL-15, IL-18, SDF-1, and TSLP.

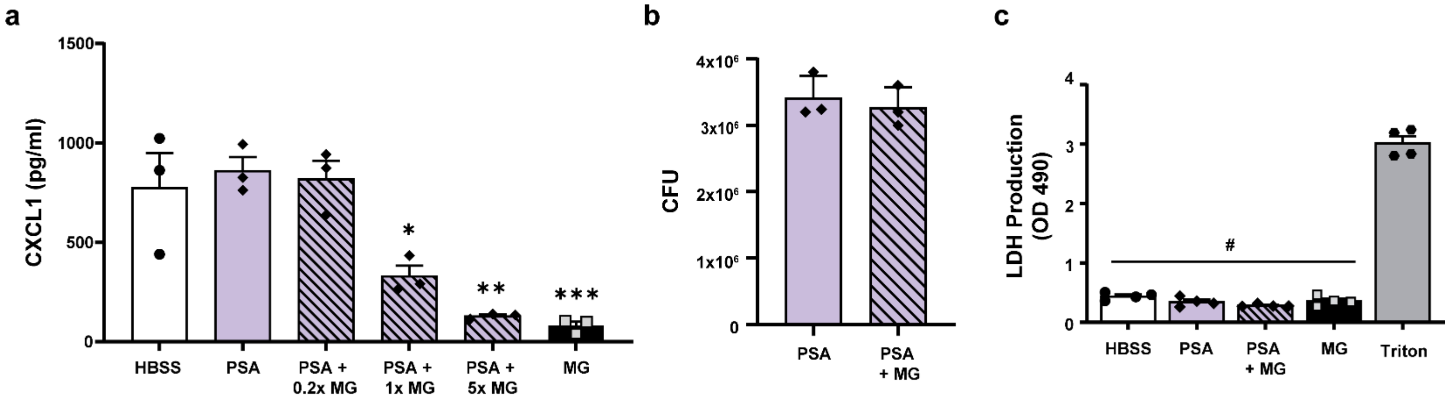


**Supplemental Figure 3. Melanin blocks *Pseudomonas*-induced CXCL1 induction and does not impact adherence or cell viability. (a)** H292 epithelium infected with media alone (HBSS; negative control), *P. aeruginosa* PAO1 strain (PSA) in the absence and presence of increasing *Aspergillus* melanin ghosts (MG; 0.2x= $2x10^6/cm^2$; 1x= $1x10^7/cm^2$; 5x= $5x10^7/cm^2$), or MG alone for 6h. CXCL1 secretion in the supernatant was measured by ELISA. n=3. One-way ANOVA with Tukey's multiple comparisons test; *p<0.05, **p<0.01, ***p<0.001 when compared to PSA alone. **(b)** Following infection with PSA ± MG ($5x10^7/cm^2$) for 6h, H292 cells were washed with media to remove any non-adherent bacteria, then cells were lysed using Triton X-100 to liberate both adhered and internalized bacteria. Serial dilutions were plated on LB agar and grown at 37°C. Colonies were counted after 24h. n=3. Unpaired t-test; no significance. **(c)** Cell viability was measured by LDH assay. H292 cells were treated with media alone, PSA ± MG ($5x10^7/cm^2$), MG alone, or 1% Triton. Treatment with Triton leads to rapid cell lysis and release of LDH, thus was used as a positive control. n=4. One-way ANOVA with Tukey's multiple comparisons test; #p<0.0001 compared to positive control. Data are represented as mean ± SEM.

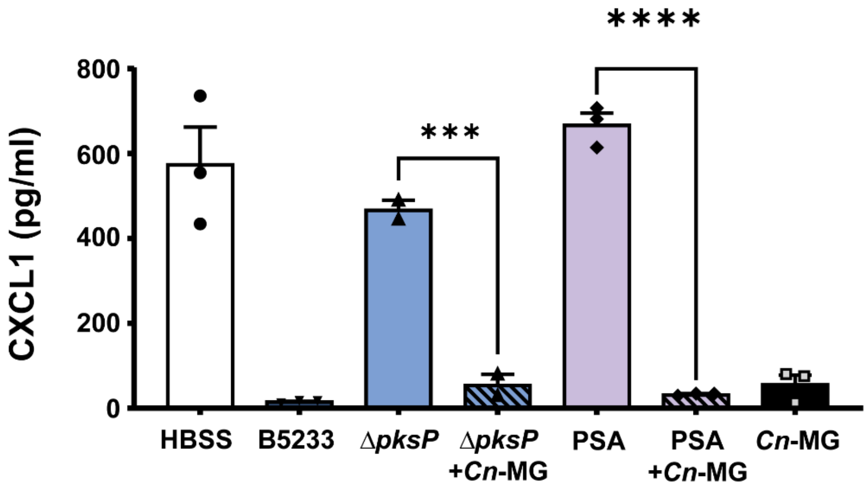


**Supplemental Figure 4.** H292 cells were stimulated for 6h with wildtype *A. fumigatus* (B5233; $1x10^7/cm^2$), *ΔpksP* conidia ($1x10^7/cm^2$) with or without *C. neoformans* melanin ghosts (*Cn*-MG; $5x10^7/cm^2$), *P. aeruginosa* PAO1 strain (PSA) with or without *Cn*-MG, or *Cn*-MG alone ($5x10^7/cm^{2)}$. CXCL1 secretion was measured by ELISA. Data are represented as mean ± SEM. n=3; One-Way ANOVA with Tukey's multiple comparisons test ***p<0.001, ****p<0.0001 compared to stimulation alone (*i.e., ΔpksP* or PSA alone).

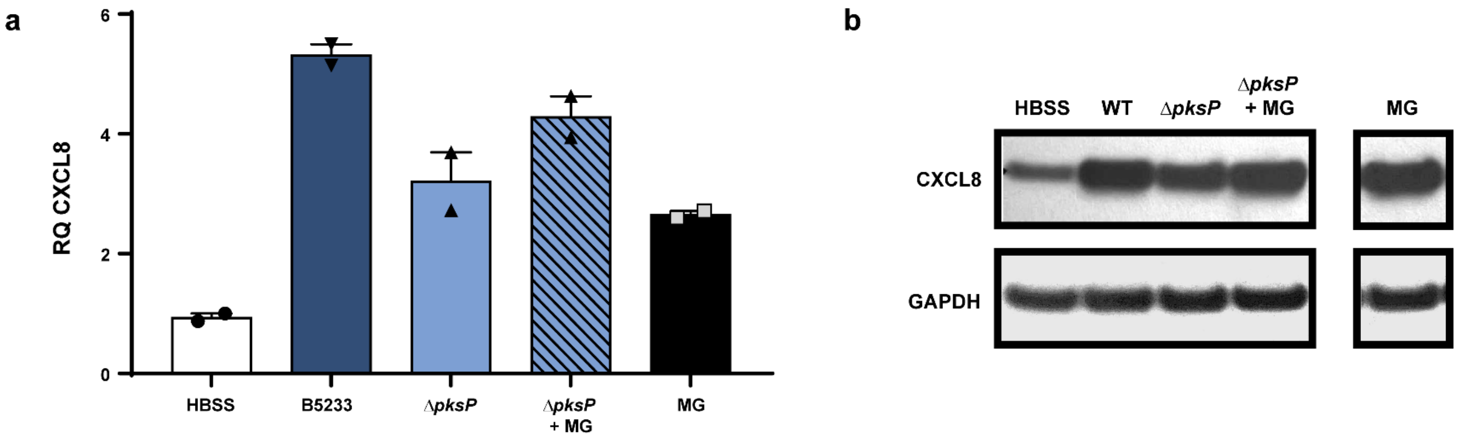


**Supplemental Figure 5.** CXCL8 RNA **(a)** and protein **(b)** expression in primary hAECs cell lysates were measured by qPCR and Western blot, respectively, following 6h stimulation with *A. fumigatus* wildtype (WT; B5233 strain; $1x10^7/cm^2$), *ΔpksP* conidia ($1x10^7/cm^2$) ± MG, *P. aeruginosa* PAO1 strain (PSA) ± MG, or MG alone ($5x10^7/cm^2$). For Western blots, GAPDH was used as a loading control. n=2. Data are represented as mean ± SEM.
